# Complete absence of thebaine biosynthesis under home-brew fermentation conditions

**DOI:** 10.1101/024299

**Authors:** Drew Endy, Stephanie Galanie, Christina D. Smolke

**Affiliations:** Department of Bioengineering; 443 Via Ortega, MC 4245 Stanford University; Stanford, CA 94305; Department of Chemistry; 443 Via Ortega, MC 4245 Stanford University; Stanford, CA 94305

## Abstract

Yeast-based biosynthesis of medicinal compounds traditionally derived from plant materials is improving. Both concerns and hopes exist for the possibility that individual small volume batch fermentations could provide distributed and independent access to a diversity of compounds some of which are now abused, illegal, or unavailable to many who need for genuine medical purposes. However, there are differences between industrial bioreactors and ‘home-brew’ fermentation. We used engineered yeast that make thebaine, a morphinan opiate, to quantify if differences in fermentation conditions impact biosynthesis yields. We used yeast that make an English ale as a positive fermentation control. We observed no production of thebaine and miniscule amounts of reticuline, an upstream biosynthetic intermediate, in home-brew fermentations; the positive control was palatable. We suggest that additional technical challenges, some of which are unknown and likely unrelated to optimized production in large-volume bioreactors, would need to be addressed for engineered yeast to ever realize home-brew biosynthesis of medicinal opiates at meaningful yields.

## Introduction

Once discovered and developed, most Western medicines are manufactured and made available via centralized and regulated industrial supply chains [Liu, 2011; World Health Organization, 2011]. For example, in 1999 almost 93 percent of global pharmaceutical production by value occurred in countries with a gross national product per capita above $9,360, with the top five producing countries (USA, Japan, France, Germany, UK) accounting for ∼67 percent of production by value [World Health Organization, 2011]. However, the majority of people who need medicines cannot reliably afford or even access them [Seya et al., 2011].

Yeast are naturally occurring microorganisms that live on every continent including Antarctica [Carrasco et al., 2012]. Humans have adapted yeast to leaven bread and brew wine or beer [Mortimer, 2000]. Fermentation with adapted yeast is widely practiced by diverse peoples, from subsistence farmers in Northern Nigeria [Netting, 1964] to citizens of modern industrialized nations who might otherwise favor specialization of labor and centralized manufacturing [Enkerli, 2006].

Following the development of recombinant DNA technology [Jackson et al., 1972] yeast have been directly engineered to make various substances, from bulk and fine chemicals to active pharmaceutical ingredients [Li & Borodina, 2014; Siddiqui et al., 2012]. In cases where yeast is used to make a product that already exists, yeast-based fermentations can displace existing supply chains. Such displacements can be disruptive. For example, yeast-based biosynthesis enabling production of semi-synthetic artemisinin is expected to both lower the price and stabilize the supply of an essential antimalarial medicine [Paddon & Keasling, 2014]. Because yeast use sugar as their primary carbon and energy source, the agricultural input to a yeast-based manufacturing process can be decoupled from the resulting product. Thus, a few crop plants (rice, corn, sugarcane, beets, potatoes) can be optimized for intensive agricultural production of commodity feedstock sugars while many different yeast strains are engineered to produce a diversity of products. As a result, land use and employment are impacted. For example, yeast-based biosynthesis of artemisinin is estimated to reduce agricultural land use and labor requirements 35-fold and 1000-fold, respectively, relative to traditional sourcing via cultivation of sweet wormwood [Jim Thomas, personal communication].

Yeast have very recently been engineered to make medicinal opioids at low titers [Galanie et al., 2015]. The existing supply chain for these essential and regulated medicines is again plant-based, starting with the farming of opium poppies [Galanie et al., 2015 and references therein]. In part due to widespread addiction and abuse of these compounds, many have imagined or expressed public concern at the prospect of yeast-based biosynthesis of opioids by individuals via home-brew fermentation. For example, Professor Voigt of MIT recently stated that “It is going to be possible to ‘home-brew’ opiates in the near future” and that a dose could be obtained from “a glass of yeast culture grown with sugar on a windowsill” [Begley, 2015]. Professor Oye of MIT argued that access to yeast strains engineered to produce narcotics should be restricted to licensed facilities, authorized researchers, and technicians [Oye et al., 2015].

However, it is not entirely obvious that restricted access to or criminalization of controlled substances leads to better public health outcomes [Greenwald, 2009]. To inform conversations and policy considerations we decided to test if yeast recently engineered to produce thebaine starting from sugar under laboratory conditions would also produce thebaine in simple home-brew fermentations [Figure 1; Galanie et al., 2015].

**Figure 1.**
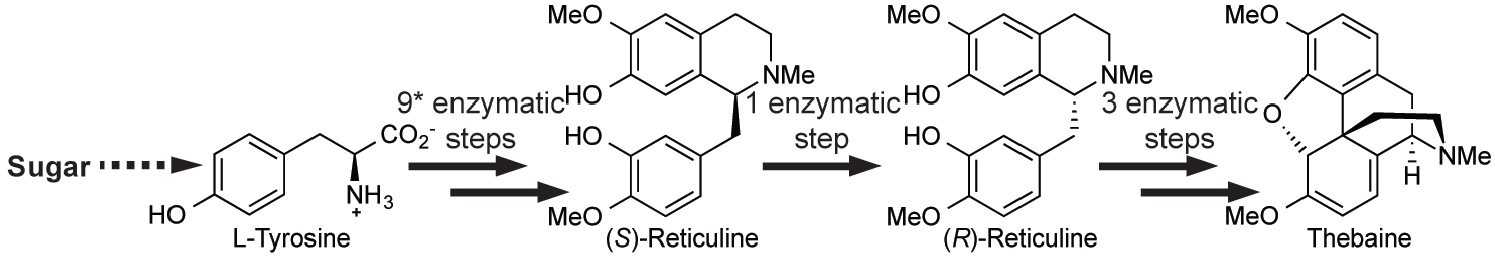
Engineered biosynthetic pathway in yeast for production of the morphinan opiate thebaine from simple carbon and nitrogen sources. * indicates that there are an additional 5 enzymes engineered into the strain for biosynthesis, recycling, and salvage of the mammalian redox cofactor tetrahydrobiopterin. See [Galanie et al., 2015] for complete strain details.

## Materials and Methods

Autoclave-sterilized glass fermentation bottles (32 ounce swing top, More Beer, Inc.) were filled with 500 mL of media. Media was 125 g/L dried malt extract (More Beer, Inc.) in water, autoclaved for 15 minutes. Single colonies of CSY1064 + pYES1L/D19CjNCS yeast, engineered to produce the morphinan opiate thebaine [Galanie et al., 2015], were inoculated into 3 mL yeast nitrogenous base media (YNB) with –Trp drop-out supplement, grown 17 h, and then used to inoculate a 50 mL culture. When the cultures reached OD_600_ 4.5, the culture (∼3E9 cells) was pelleted by centrifugation and resuspended in 1 mL sterile water. The fermentation bottles were inoculated with this resuspended yeast or with, as a positive fermentation control, 390 mg Safale S-04 yeast (∼3E9 cells, More Beer, Inc.). The fermentation bottles were sealed with a #2 stopper and 3-piece airlock (More Beer, Inc.), and stored in a secure, roomtemperature environment [Figure 2]. After 120 h, 1 mL samples were removed, centrifuged 10 min at full speed to precipitate yeast and particulates, and analyzed to determine reticuline and thebaine concentrations by high performance liquid chromatography-tandem mass spectrometry (HPLC-MS/MS) using multiple reaction monitoring (MRM) according to our previously published method [Galanie et al., 2015]. The Safale S-04 positive fermentation control was also tested by tasting.

**Figure 2.**
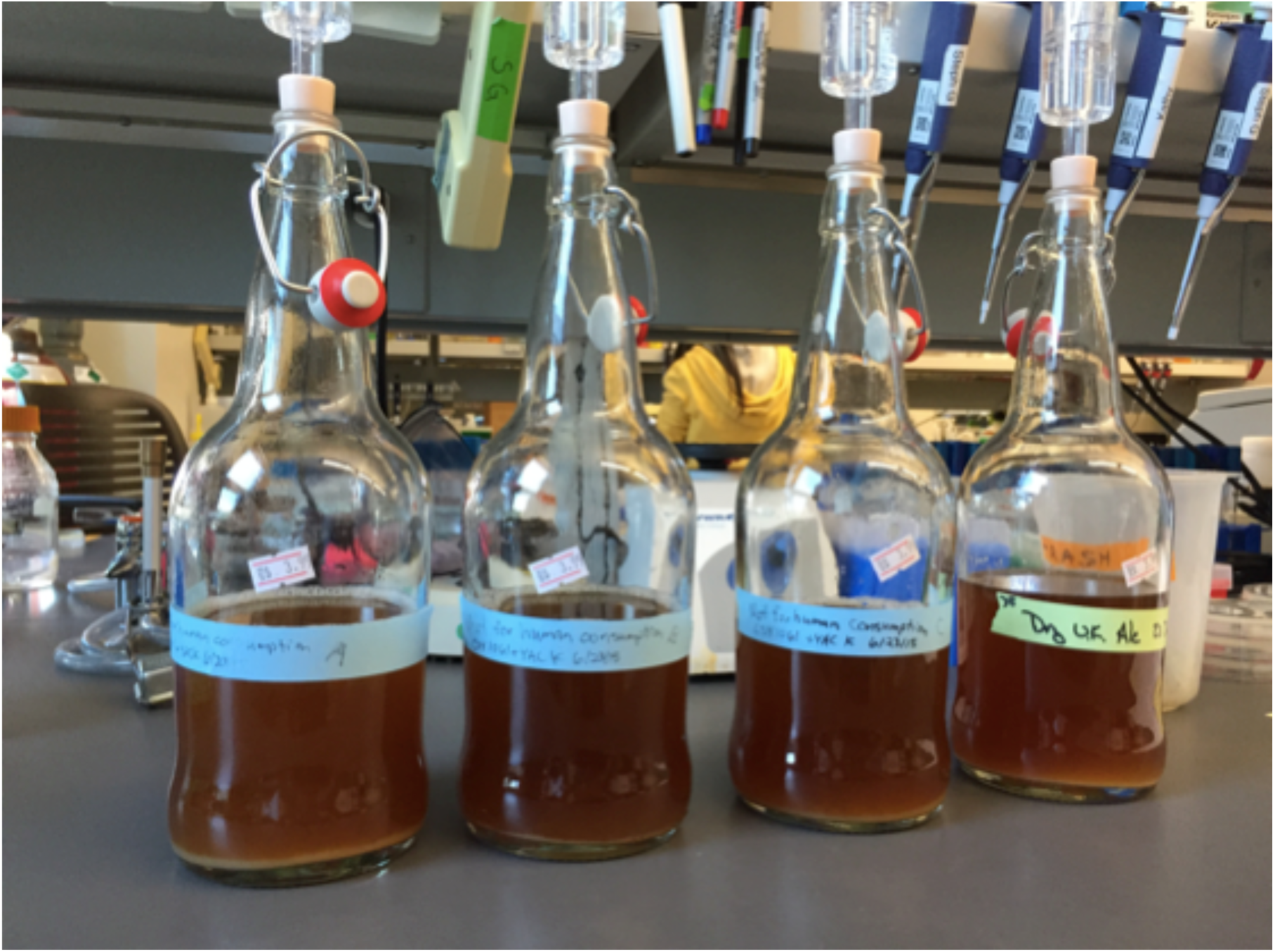
Small volume ‘home-brew’-style fermentation of yeast engineered to produce thebaine. (three left bottles) **or adapted to produce an English ale** (right bottle, positive fermentation control).

## Results and Discussion

After culturing yeast engineered to produce the opioid drug precursor thebaine and the brewing strain control Safale S-04 under non-laboratory fermentation conditions for 120 h, we analyzed the culture media by HPLC-MS/MS. No thebaine was detected for either strain, and a trace amount (<3 ug/L) of reticuline was detected for strain CSY1064+ pYES1L/D19CjNCS but not for the Safale strain [Figure 3]. These results are markedly different from those obtained under laboratory fermentation conditions, in which a similar yeast strain produced 31.3 ug/L reticuline and 6.4 ug/L thebaine [Galanie et al., 2015]. Thus, under non-laboratory fermentation conditions yeast produced less than one-tenth of a key morphinan alkaloid precursor compared to laboratory conditions, and no detectable morphinan alkaloids.

**Figure 3.**
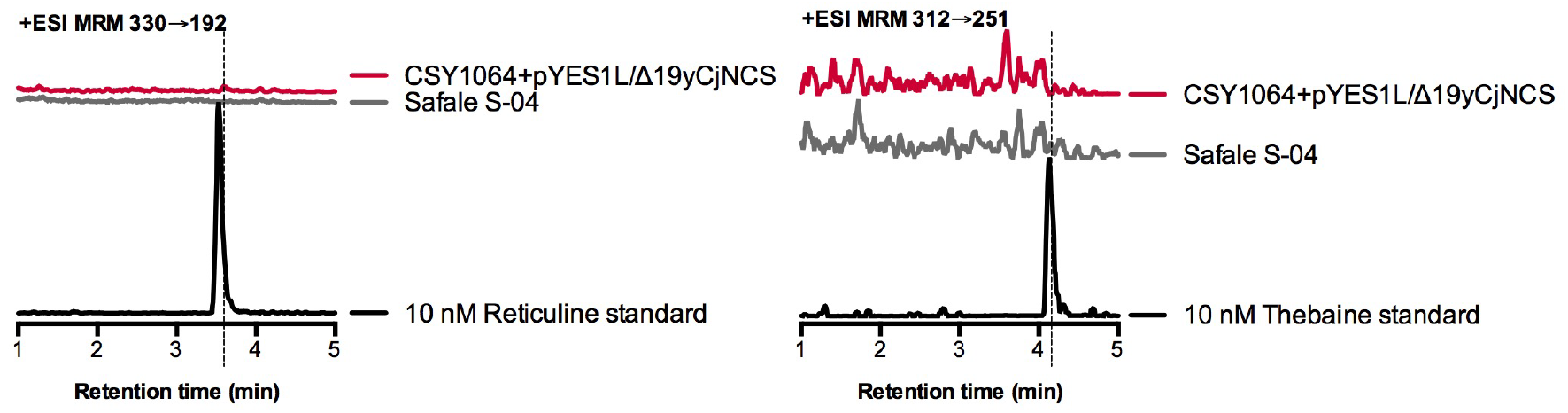
Yeast do not produce thebaine and only miniscule amounts of reticuline in home brew fermentations. Liquid chromatography-tandem mass spectrometry analysis of fermentation broth. Chromatogram traces of reticuline and thebaine in growth media for indicated strains. Traces for CSY1064+pYES1L/D19CjNCS are representative of three biological replicates. Thebaine was not detected and reticuline was detected at just above the limit of detection.

## Conclusions

The first example of yeast engineered to produce opioids from sugar under laboratory conditions does not produce detectable amounts of natural opiate or semi-synthetic opioid drug molecules in simple home-brew fermentations. Future yeast strains optimized for improved yields under laboratory conditions or in industrial fermentors might also be expected to have greatly reduced product yields in home-brew fermentations. We suggest that researchers carrying out work to improve biosynthesis yields of controlled substances also check to see how future strains perform under non-laboratory conditions and, if warranted, engineer strains that do not produce controlled substances in uncontrolled environments. We additionally support open discussion of strategies and goals for the development of microbial biosynthesis of active pharmaceutical compounds. Such discussions should include researchers, policy experts, regulatory and enforcement officials, health and medical professionals, and representatives of communities in which essential medicines are either unavailable or abused.

